# The type IVa pilus machinery is pre-installed during cell division

**DOI:** 10.1101/087965

**Authors:** Tyson Carter, Ryan N.C. Buensuceso, Stephanie Tammam, Ryan P. Lamers, Hanjeong Harvey, P. Lynne Howell, Lori L. Burrows

**Affiliations:** Department of Biochemistry and Biomedical Sciences and the Michael G. DeGroote Institute for Infectious Disease Research, McMaster University, ON, CANADA; Program in Molecular Structure & Function, The Hospital for Sick Children; Department of Biochemistry, University of Toronto, ON, CANADA

**Author notes:** To whom correspondence should be addressed: Dr. Lori L. Burrows, 4H18 Health Sciences Centre, 1200 Main St. West, Hamilton, ON, L8N 3Z5 CANADA, Dr. P. Lynne Howell, 20-9-715 Peter Giligan Centre for Research and Learning, 686 Bay St., Toronto, ON M5G 0A4 CANADA.

## Abstract

Type IV pili (T4aP) are ubiquitous microbial appendages used for adherence, twitching motility, DNA uptake, and electron transfer. Many of these functions depend on dynamic assembly and disassembly of the pilus by a megadalton-sized, cell envelope-spanning protein complex located at the poles of rod-shaped bacteria. How the T4aP assembly complex becomes integrated into the cell envelope in the absence of dedicated peptidoglycan (PG) hydrolases is unknown. After ruling out potential involvement of housekeeping PG hydrolases in installation of the T4aP machinery in *P. aeruginosa*, we discovered that key components of inner (PilMNOP) and outer (PilQ) membrane subcomplexes are recruited to future sites of cell division. Mid-cell recruitment of a fluorescently tagged alignment subcomplex component, mCherry-PilO, depended on PilQ secretin monomers – specifically, their N-terminal PG-binding AMIN domains. PilP, which connects PilO to PilQ, was required for recruitment, while PilM, which is structurally similar to divisome component FtsA, was not. Recruitment preceded secretin oligomerization in the outer membrane, as loss of the PilQ pilotin, PilF, had no effect on localization. These results were confirmed in cells chemically blocked for cell division prior to outer membrane invagination. The hub protein FimV and a component of the Polar Organelle Coordinator complex – PocA – were independently required for mid-cell recruitment of PilO and PilQ. Together, these data reveal an integrated, energy-efficient strategy for the targeting and pre-installation – rather than retrofit – of the T4aP system into nascent poles, without the need for dedicated PG-remodelling enzymes.

## IMPORTANCE

The peptidoglycan (PG) layer of bacterial cell envelopes has limited porosity, representing a physical barrier to insertion of large protein complexes involved in secretion and motility. Many systems include dedicated PG hydrolase components that create space for their insertion, but the ubiquitous type IVa pilus (T4aP) system lacks such an enzyme. Instead, we found that components of the T4aP system are recruited to future sites of cell division where they can be incorporated into the cell envelope during the formation of new poles, eliminating the need for PG hydrolases. Targeting depends on the presence of septal PG-binding motifs in specific components, as removal of those motifs causes delocalization. This pre-installation strategy for the T4aP assembly system ensures that both daughter cells are poised to extrude pili from new poles as soon as they separate from one another.

## INTRODUCTION

The cell envelopes of most gram-negative bacteria are comprised of an inner membrane, a peptidoglycan (PG) layer found in the periplasm, and an outer membrane (OM). A variety of large protein complexes, including motility machines, secretion systems, and efflux pumps span all these layers (1), but in many cases the mechanisms used to integrate these systems into the cell envelope remain uncharacterized. In many cases, they are integrated at specific locations in the cell such as the poles, which is important for their function.

The machinery that assembles and disassembles type IVa pili (T4aP) is among the most common cell envelope-spanning complexes found in gram-negative bacteria. T4aP are long, thin protein fibres used for adherence, biofilm formation and a flagellum-independent form of motility known as twitching, defined as repeated cycles of pilus extension, adherence to a surface, and retraction of the pilus fibre (2, 3). *P. aeruginosa* T4aP are important for surface mechanosensing and consequent up-regulation of virulence factor expression (4), and for the delivery of toxins by other secretion systems (5, 6). In the absence of T4aP, *P. aeruginosa* has reduced pathogenicity (5).

The T4aP machinery is composed of four subcomplexes that together form a dynamic cylindrical machine (3, 7, 8). The inner membrane motor and alignment subcomplexes form the platform for assembly and disassembly of the pilus, and interact directly with the secretin that allows for pilus extrusion through the OM. The OM lipoprotein PilF promotes formation of the secretin, a highly stable oligomer of 14 PilQ subunits that form a gated channel in the OM (9, 10). The N-terminus of each PilQ monomer contains two tandem amidase N-terminal (AMIN) domains, involved in binding peptidoglycan (11–13). The alignment subcomplex composed of PilMNOP connects the motor subcomplex with the secretin, and participates in pilus extension and retraction (14–17). PilM is a cytoplasmic, actin-like protein that clamps tightly onto the short amino-terminus of the inner membrane protein, PilN, which in turn forms heterodimers with inner membrane protein, PilO (14, 18, 19). The inner membrane lipoprotein PilP interacts with PilNO heterodimers via its long unstructured N-terminus, and with the periplasmic N0 domain of PilQ via its C-terminal homology region (HR) domain, completing the transenvelope complex (17, 20).

T4aP are usually located at the poles of rod-shaped cells, which promotes adherence by minimizing surface area and thus electrostatic repulsion (3). How the multi-protein T4aP system is initially targeted to and integrated into the poles of *P. aeruginosa* cells are not well understood. Type III (T3SS) and Type IV (T4SS) secretion systems have dedicated PG-remodelling enzymes that allow for retrofitting of the complexes into the cell envelope (1). There are no such enzymes associated with the T4aP system, suggesting that it uses an alternate pathway for cell envelope integration. Here we used fluorescent fusions to an informative alignment subcomplex component, PilO, and to the secretin monomer, PilQ, to track and quantify their localization in *P. aeruginosa* wild type and mutant backgrounds. The results show that PilO and associated components are recruited to future sites of cell division by the concerted action of least three pathways. We propose that the T4aP machinery is preinstalled during formation of nascent cell poles, a strategy that eliminates the need for dedicated cell-wall-processing enzymes.

## RESULTS

### mCherry-PilO localizes to cell poles and future cell division sites

When selecting components of the T4aP machinery to track using fluorescent fusions, we considered the known interactions and stoichiometry of these proteins and how a fusion might affect function. PilM is cytoplasmic, but has multiple interaction partners (14, 19, 20), including PilN, whose cytoplasmic N-terminus binds to a groove on PilM; thus, we excluded both as candidates. PilN interacts along most of its remaining length with PilO (15, 21), but PilO’s cytoplasmic N-terminus is poorly conserved among T4aP-expressing species (16), supporting the observation that it appears to have no cytoplasmic interaction partners. Therefore, PilO was selected as a fusion candidate for localization of the alignment subcomplex. To maintain the physiological expression levels and stoichiometry that are important for function (14), a *pilO* construct encoding an in-frame fusion of mCherry to the N-terminus of PilO was used to replace the WT version at the native *pilO* locus (**Fig. 1A**). Stability and functionality of the fusion protein were verified by western blot with anti-PilO and anti-mCherry antisera, and assessment of twitching motility, respectively (**Fig. 1B**).

**Figure 1.**
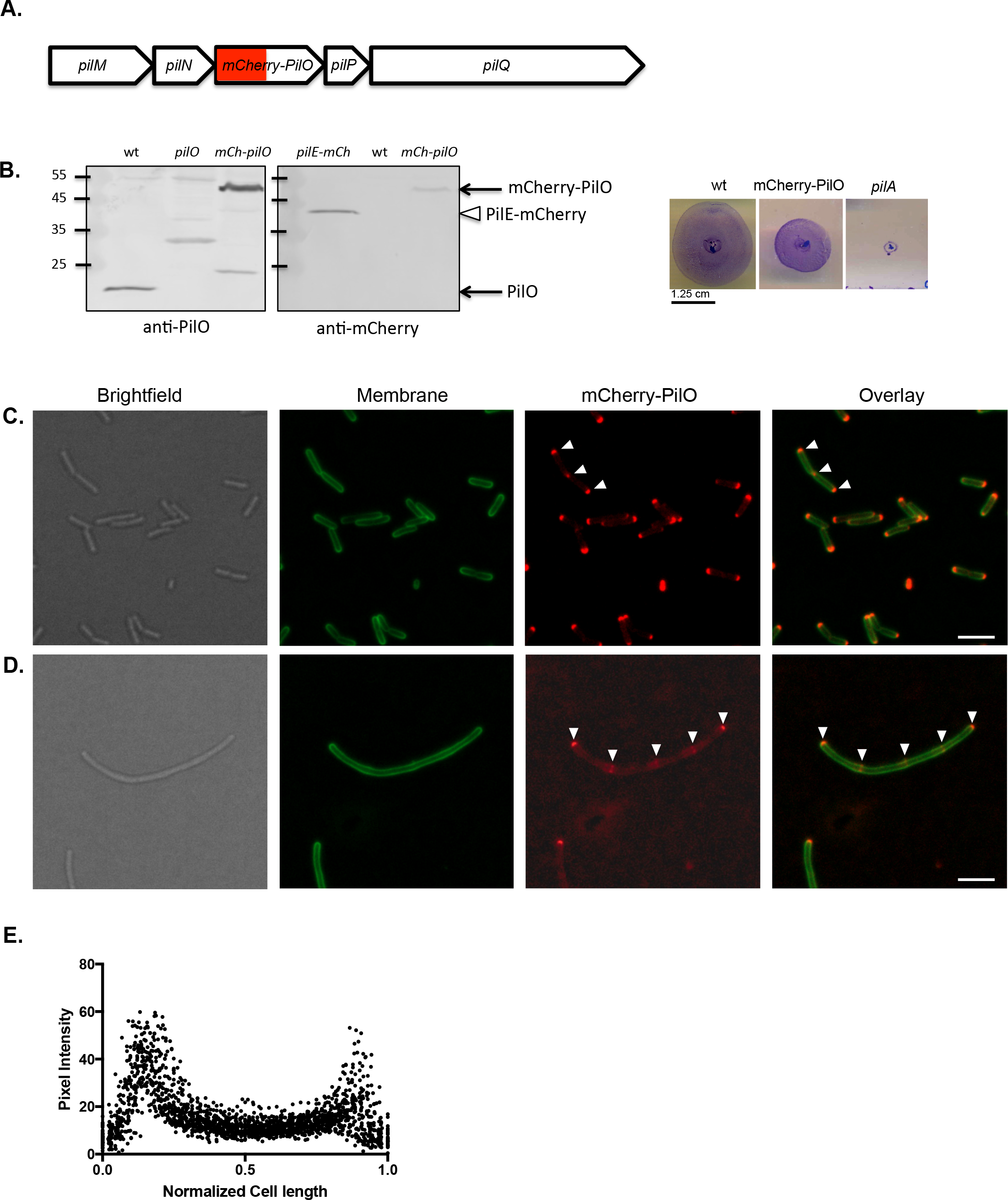
mCherry-PilO localizes to cell poles and future sites of cell division in *P. aeruginosa.* **A**. Map of the T4aP alignment subcomplex operon showing *mCherry-pilO* integration. **B**. The mCherry-PilO fusion was stable (∼48 kDa product recognized by both anti-PilO and anti-mCherry antibodies; a plasmid-encoded PilE-mCherry fusion – open triangle – was used as a positive control for the latter; (64)) and functional for twitching motility. **C**. mCherry-PilO localized to the poles of wild type cells and to the septum in late-stage dividing cells. **D**. When cells were filamented using 40 μg/ml cefsulodin, mCherry-PilO localized to the poles and at regularly spaced foci (arrowheads). Scale bar = 3 μm. **E**. mCherry pixel intensity in 50 untreated cells after normalization of length to 1 as described in the Methods.

mCherry-PilO localized to both poles, as well as to the midpoint of cells undergoing division (**Fig. 1C**). To more clearly visualize the predivisional localization pattern of mCherry-PilO, cells were treated with sub-inhibitory levels of cefsulodin, which inhibits PBP3 (FtsI), a late-stage PG transpeptidase, leading to division arrest and filamentation (22). In filamented cells, mCherry-PilO localized to the poles and to regularly spaced foci, suggestive of recruitment to future sites of cell division (**Fig. 1D**). Quantification of mCherry-PilO localization in wild type cells (**Fig. 1E**) confirmed its predominately bipolar localization.

To provide further evidence that mCherry-PilO was recruited to sites of cell division, the *minC* gene, which encodes a key component of the oscillating Min system required for inhibition of FtsZ polymerization at non-mid-cell locations (23, 24) was deleted in the mCherry-PilO strain. Cells lacking MinC exhibit aberrantly localized septa, producing mini-cells due to unequal cell division (**Fig. 2A**). As predicted, the mCherry-PilO fusion continued to localize to polar and septal sites despite their aberrant placement (**Fig. 2B**). Together, these data suggest that when expressed under physiological conditions, PilO – and by inference, its interaction partners, PilMN and PilP (20) – are recruited to future sites of cell division.

**Figure 2.**
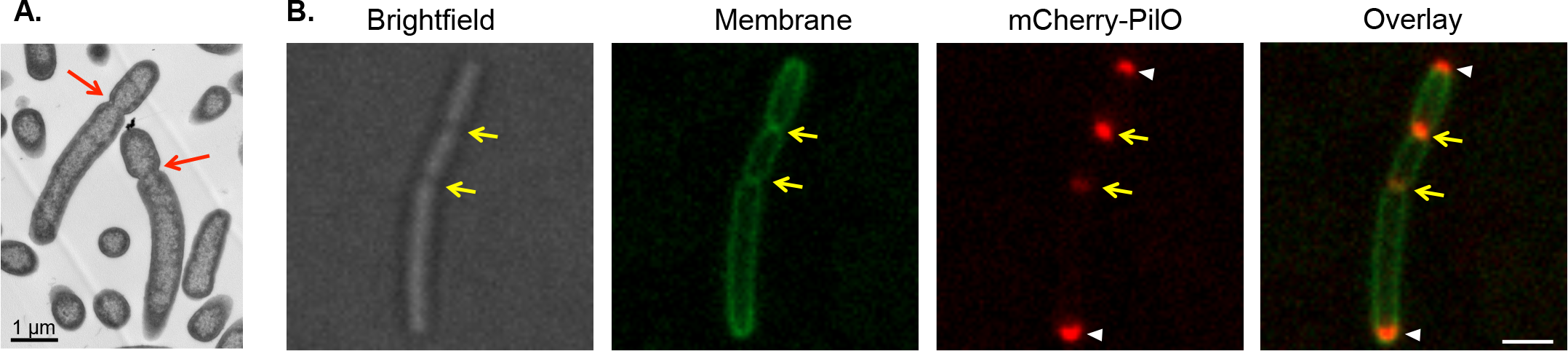
mCherry-PilO localizes to aberrantly localized septa in a *minC* mutant. **A**. Transmission electron micrograph of a *minC* mutant of *P. aeruginosa* strain PAK. In the absence of *minC*, cells have aberrantly located division sites (red arrows). Scale bar = 1 μm. **B**. mCherry-PilO polar foci are indicated with white arrowheads while foci localized to aberrantly placed septa are indicated with yellow arrows.

### PilQ monomers are required for T4aP alignment subcomplex localization

To define the mechanism of recruitment of the T4aP alignment subcomplex to sites of cell division, we first examined the effects of deleting specific subcomplex components. The cytoplasmic member of the alignment subcomplex, PilM, has pronounced structural similarity (Cα root mean square deviation = 1.9 Å over 257 residues) to the early divisome protein, FtsA, a peripheral inner membrane protein whose interaction with FtsZ tethers the latter to the membrane (18, 19). Based on its structural mimicry of FtsA and its peripheral membrane localization via binding of PilN’s amino terminus, we previously hypothesized (25) that PilM – and thus the alignment subcomplex – might be recruited to mid-cell through interactions with the FtsZ ring. However, mCherry-PilO exhibited polar and mid-cell localization in a *pilM* mutant (**Fig. 3AB**), ruling out PilM as a driver of recruitment.

**Figure 3.**
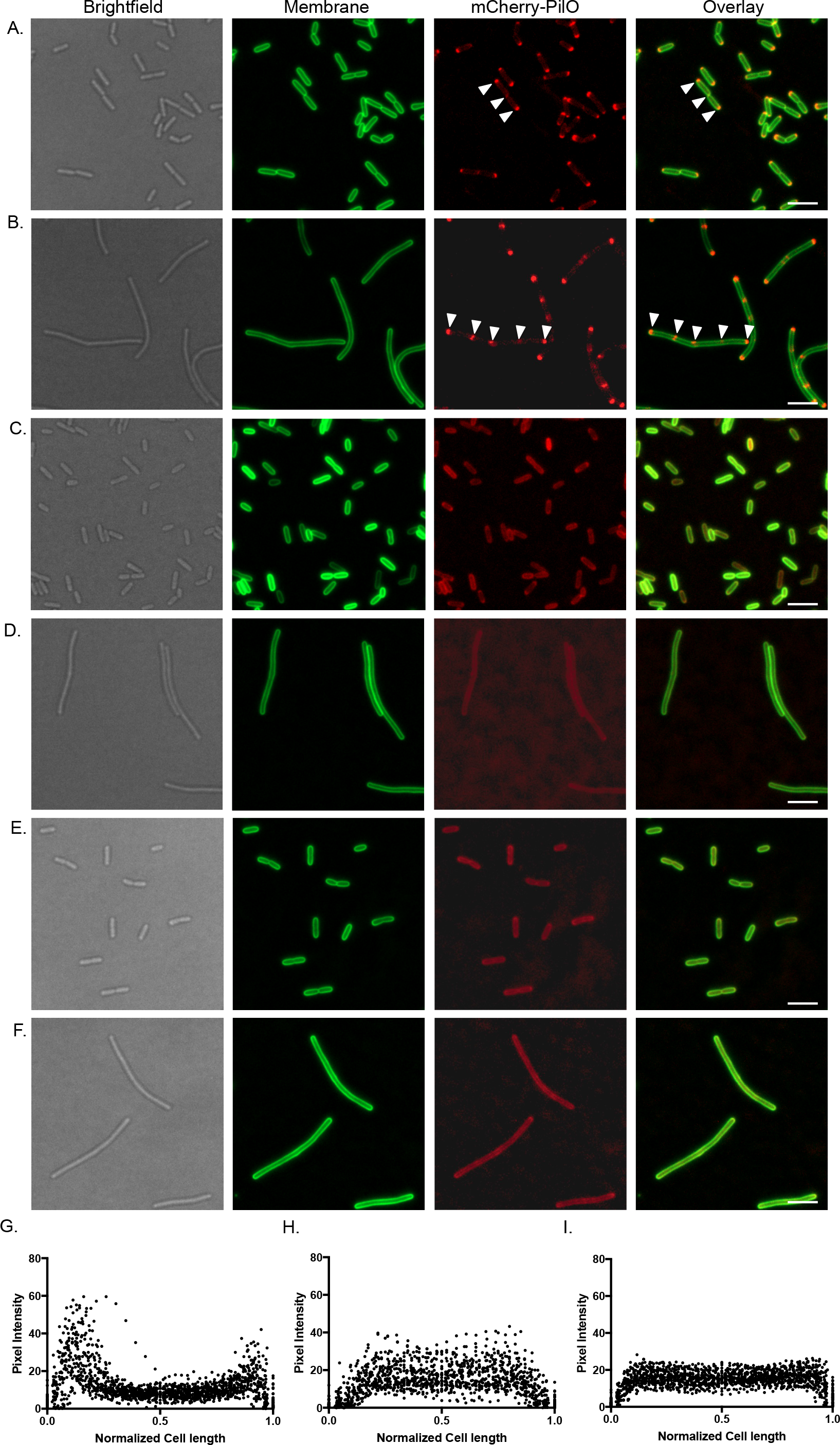
Polar localization of T4aP alignment subcomplex requires PilQ. In the absence of PilM, mCherry-PilO is localized to **A**. poles and septa in untreated cells, and poles and regularly spaced foci in cells treated with cefsulodin. In the absence of PilQ, mCherry-PilO is delocalized in both **C**. untreated and **D**. antibiotic-filamented cells. Similarly, in the absence of PilP, mCherry-PilO is delocalized in **E**. untreated and **F**. antibiotic-treated cells. The last 3 panels show quantification of mCherry pixel intensity in the absence of **G**. *pilM*, **H**. *pilQ*, or **I**. *pilP*, in 50 untreated cells after normalization of length to 1 as described in the Methods. Scale bar = 3 μm.

The PilMNOP alignment subcomplex connects to the secretin via interaction of the C-terminal HR domain of PilP with the periplasmic N0 domain of PilQ (20). Preceding the N0 domain in *P. aeruginosa* PilQ are two tandem AMIN domains, predicted to bind septal PG (11–13). To test their PG-binding ability, we cloned, expressed, and purified 3 fragments of PilQ’s N-terminal region (the AMIN domains alone; the N0-N1 domains alone; and all 4 domains together), and performed pulldown assays using purified *P. aeruginosa* PG (**Supplementary Fig. S1**). In the absence of PG, all fragments remained in the soluble fraction; however, when insoluble PG was present, only those fragments containing the AMIN domains were also found in the insoluble fraction, indicating that they bind PG.

Since *pilMNOPQ* are expressed from a single operon, we hypothesized that recruitment of PilMNOP to sites of cell division might occur through a series of PilMNOP-PilQ-septal PG interactions. We first confirmed that PilQ-mCherry was localized to the poles and to regularly spaced foci in filamented cells, a pattern similar to mCherry-PilO (**Supplementary Fig. S2**). When we deleted *pilQ*, mCherry-PilO became delocalized (**Fig. 3CD**). To confirm that loss of mCherry-PilO localization in the *pilQ* strain was indirectly due to loss of PilP-PilQ interactions, *pilP* was deleted in the mCherry-PilO strain. In the *pilP* mutant, mCherry-PilO was delocalized (**Fig. 3EF**), supporting the hypothesis that recruitment of PilQ to mid-cell is the primary event, and the remaining alignment subcomplex components are recruited via their interaction with PilP, and its interaction with PilQ.

To test whether its AMIN domains were responsible for localization of PilQ and its interaction partners to mid-cell and polar positions, we complemented a *pilQ* mutant expressing mCherry-PilO with full length *pilQ*, or a construct expressing only the N0-N1 and secretin domains, previously shown to form stable but non-functional secretin oligomers (**Fig. 4A**) (9). Full length PilQ restored the wild type pattern of mCherry-PilO localization (**Fig. 4BC**), but the AMIN-deficient PilQ did not (**Fig. 4DE**). These data show that localization of T4aP assembly components is dependent upon PilQ’s AMIN domains, which interact with PG (**Supplementary Fig. S1**).

**Figure 4.**
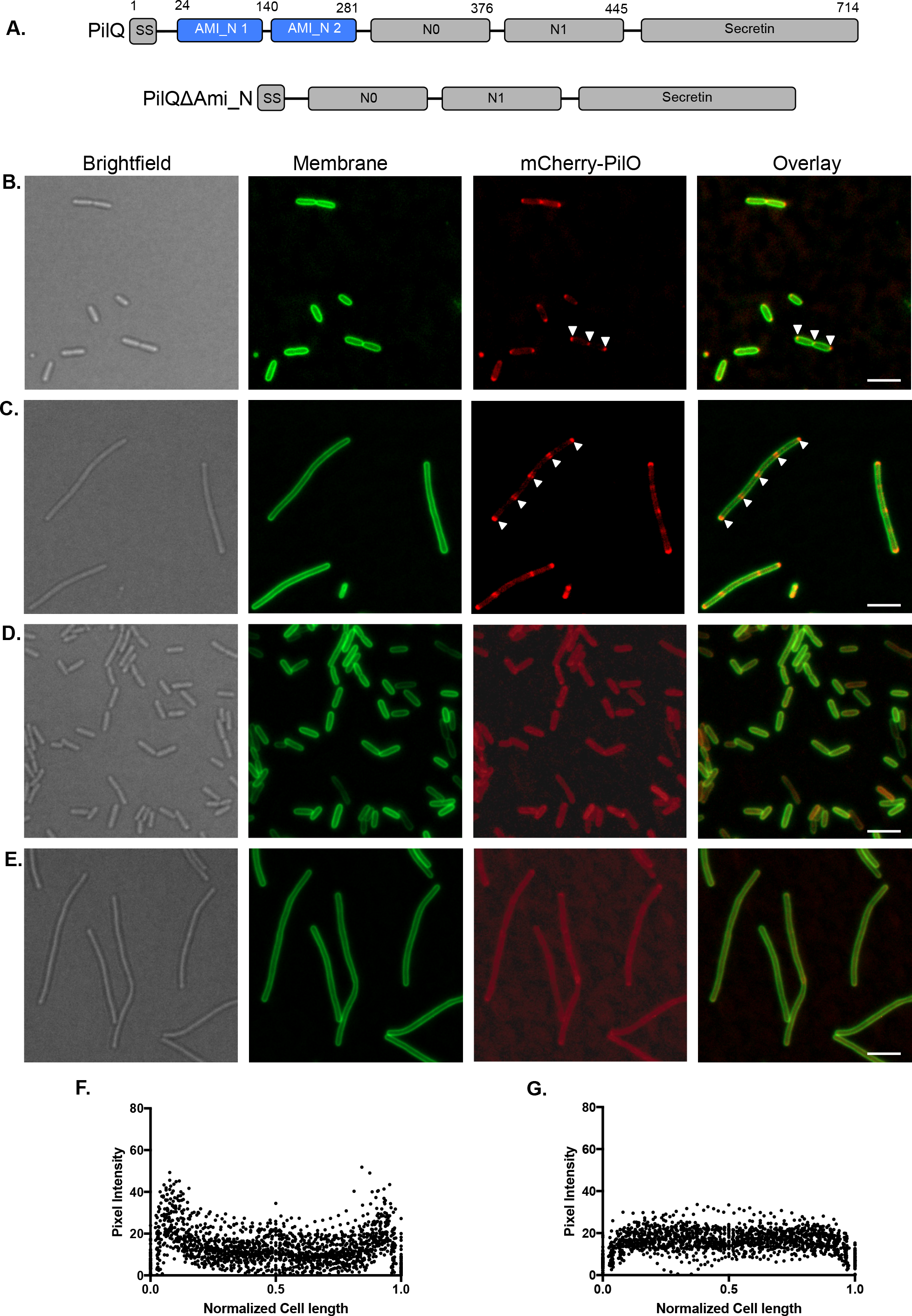
The AMIN domains of PilQ are required for polar localization of mCherry-PilO. **A**. Domain map of PilQ showing its PG-binding AMIN domains in blue. Complementation of a *pilQ* rmutant expressing mCherry-PilO with full length *pilQ* reestablished its polar and septal localization (white arrowheads) in **B**. untreated cells, and **B**. localization to the poles and regularly spaced foci in cefsulodin treated cells. When the same mutant was complemented with a truncated version of *pilQ* that encodes only N0-N1 and the secretin domains, mCherry-PilO was delocalized in both **D**. untreated and **E**. antibiotic-treated cells. The last 3 panels show quantification of mCherry pixel intensity in the presence of **F**. PilQ or **G**. PilQ lacking its AMIN domains, in 50 untreated cells after normalization of length to 1 as described in the Methods. Scale bar = 3 μm.

### Localization of mCherry-PilO precedes PilQ oligomerization

The *Myxococcus xanthus* T4aP system was proposed to assemble in an ‘outside-in’ manner, with PilMNOP alignment subcomplexes docking onto the PilQ secretin after its assembly in the OM (26). Those data are not consistent with our observation that mCherry-PilO localized to regularly spaced foci in chemically-filamented *P. aeruginosa* cells (**Fig. 1**), where the OM is not yet invaginated due to arrest of cell division. To test whether secretin oligomerization was a necessary prerequisite for recruitment, we examined localization of PilQ-mCherry in mutants lacking the OM lipoprotein PilF. In the absence of PilF, PilQ remains in a monomeric state in the inner membrane (9), but its localization pattern was unchanged compared to wild type (**Supplementary Fig. S2AB**). Similarly, localization of mCherry-PilO in the *pilF* background resembled wild type (**Fig. 5**). These data confirm that localization of T4aP components to mid-cell – while PilQ-dependent – precedes secretin oligomerization.

**Figure 5.**
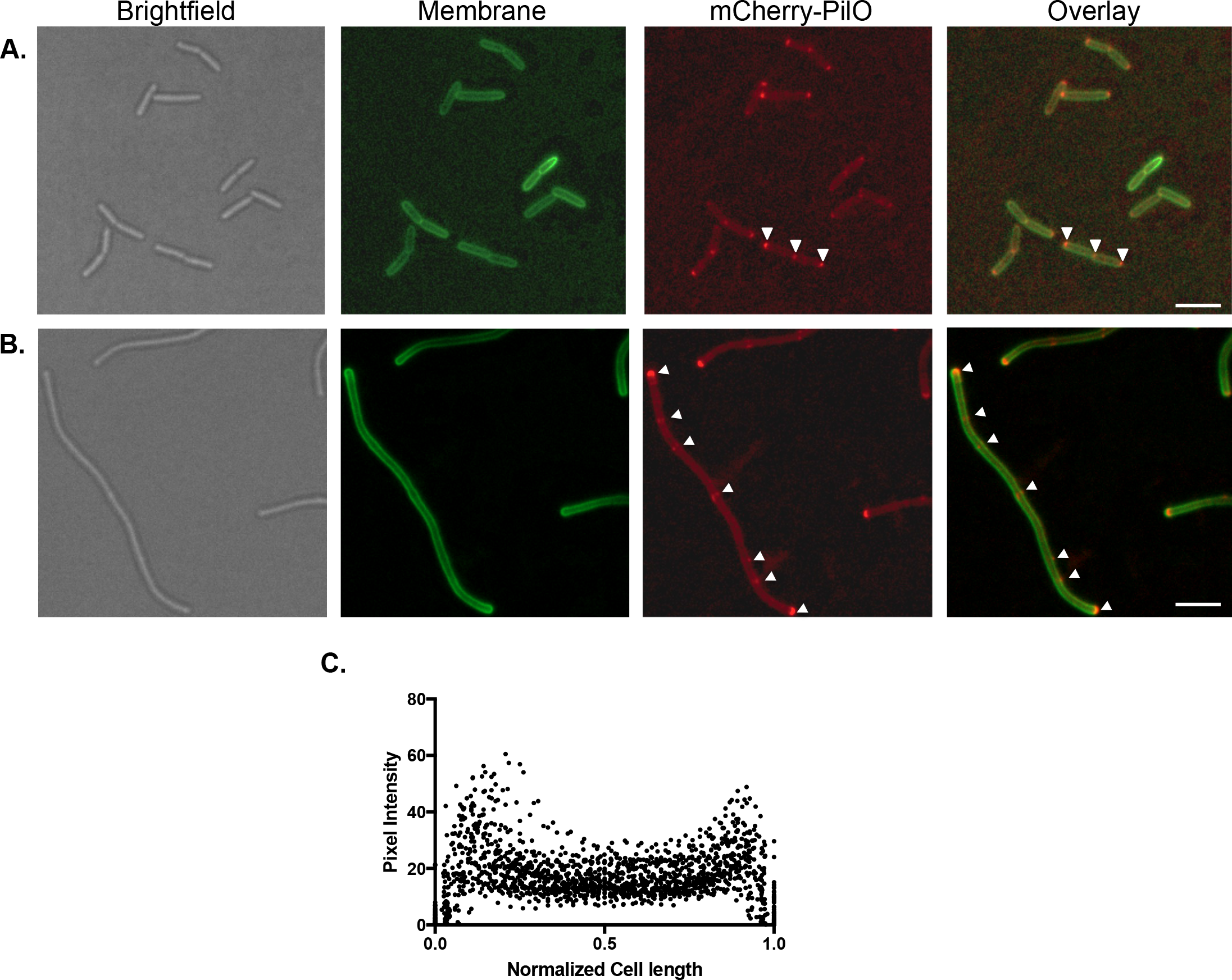
mCherry-PilO is polarly localized in the absence of *pilF*. **A**. mCherry-PilO localized to the poles and septa of late-stage dividing cells, marked by white arrowheads. **B**. When cells were filamented with cefsulodin, mCherry-PilO localized to the poles and to foci (arrows). Scale bar = 3 μm. Panel **C**. shows quantification of mCherry pixel intensity in the *pilF* background, in 50 untreated cells after normalization of length to 1 as described in the Methods.

### Additional factors modulate T4aP assembly machinery localization

FimV is a T4aP-associated protein required for twitching motility and has a highly conserved PG-binding LysM motif in its periplasmic N-terminal domain (27, 28). We showed previously (28) that PilQ multimer formation is reduced in a *fimV* mutant, which made FimV of interest for this study. In the absence of FimV, mCherry-PilO was delocalized (**Fig. 6AB**), as was PilQ-mCherry (**Supplementary Fig. S2C**). We previously generated a strain expressing a variant of FimV with an in-frame deletion of its LysM motif (28). mCherry-PilO (**Fig. 6CD**) was delocalized in the *fimV*_ΔLysM_ strain, as was the majority of PilQ-mCherry (**Supplementary Fig. S2D**). These data suggest that interactions of both PilQ and FimV with PG are important steps in the localization process, and that polar localization of PilQ depends in part on FimV.

**Figure 6.**
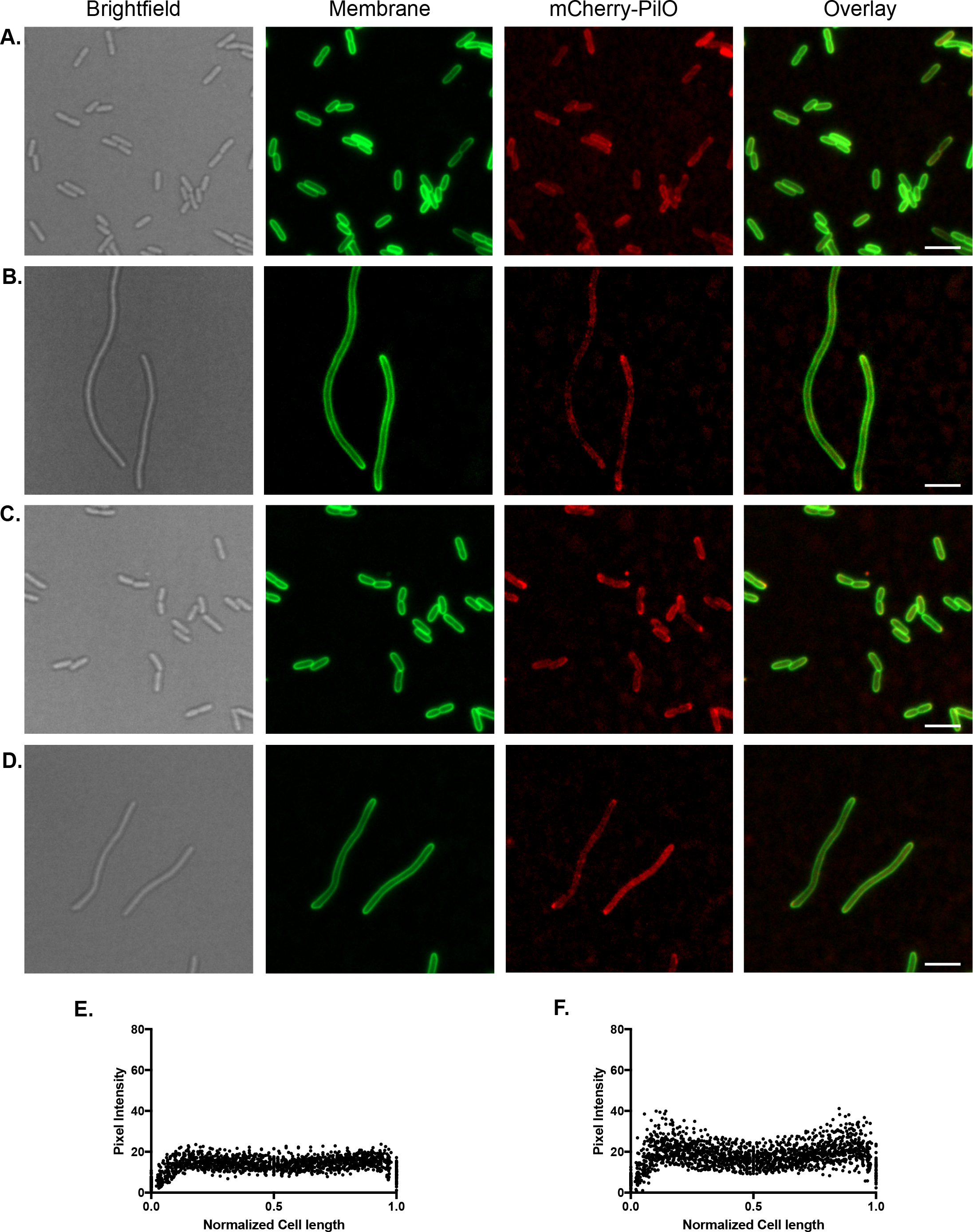
FimV is required for polar localization of mCherry-PilO. In the absence of FimV, mCherry-PilO was delocalized in **A**. untreated cells and **B**. cells treated with cefsulodin. In cells expressing a version of FimV with an in-frame deletion of its PG-binding domain (LysM), mCherry-PilO was delocalized in **C**. untreated and **D**. antibiotic-treated cells. The last two panels show quantification of mCherry pixel intensity **F**. in the absence of FimV or **G**. in the strain expressing FimV_ΔLysM_, in 50 untreated cells after normalization of length to 1 as described in the Methods. Scale Bar = 3 μm.

During a mutant screen looking for factors required for correct localization of *P. aeruginosa’s* single polar flagellum, Cowles and colleagues (29) identified a putative protein complex that they named PocAB-TonB3 for ‘polar organelle coordinator’. Loss of TonB3 had previously been shown to affect twitching motility (30), but the mechanism was unknown. All three components were required for expression and/or polar localization of T4aP. *pocA* mutants made a few, non-polar pili, while *pocB* and *tonB3* mutants failed to produce pili. The mechanism by which this system controls polar localization of motility organelles remains unclear, since the Poc proteins themselves are not localized to the poles (29).

We focused on *pocA* as the mutant’s ability to make a few pili suggested that the T4aP assembly system was functional, and deleted it from cells expressing either mCherry-PilO (**Fig. 7AB**) or PilQ-mCherry (**Supplementary Fig. S2E**). mCherry-PilO was delocalized in the absence of *pocA*, and its wild type localization was re-established upon complementation with *pocA in trans* (**Fig. 7CD**). PilQ-mCherry was delocalized in the *pocA* background although some polar fluorescence remained (**Supplementary Fig. S2E**). Interestingly, while both PocA and FimV were required for T4P assembly component localization to polar and midcell locations, they appear to act independently of one another. The fluorescent FimV fusion, which exhibits bipolar localization in wild type cells, remained localized to the poles of a *pocA* mutant (**Fig. 8**).

**Figure 7.**
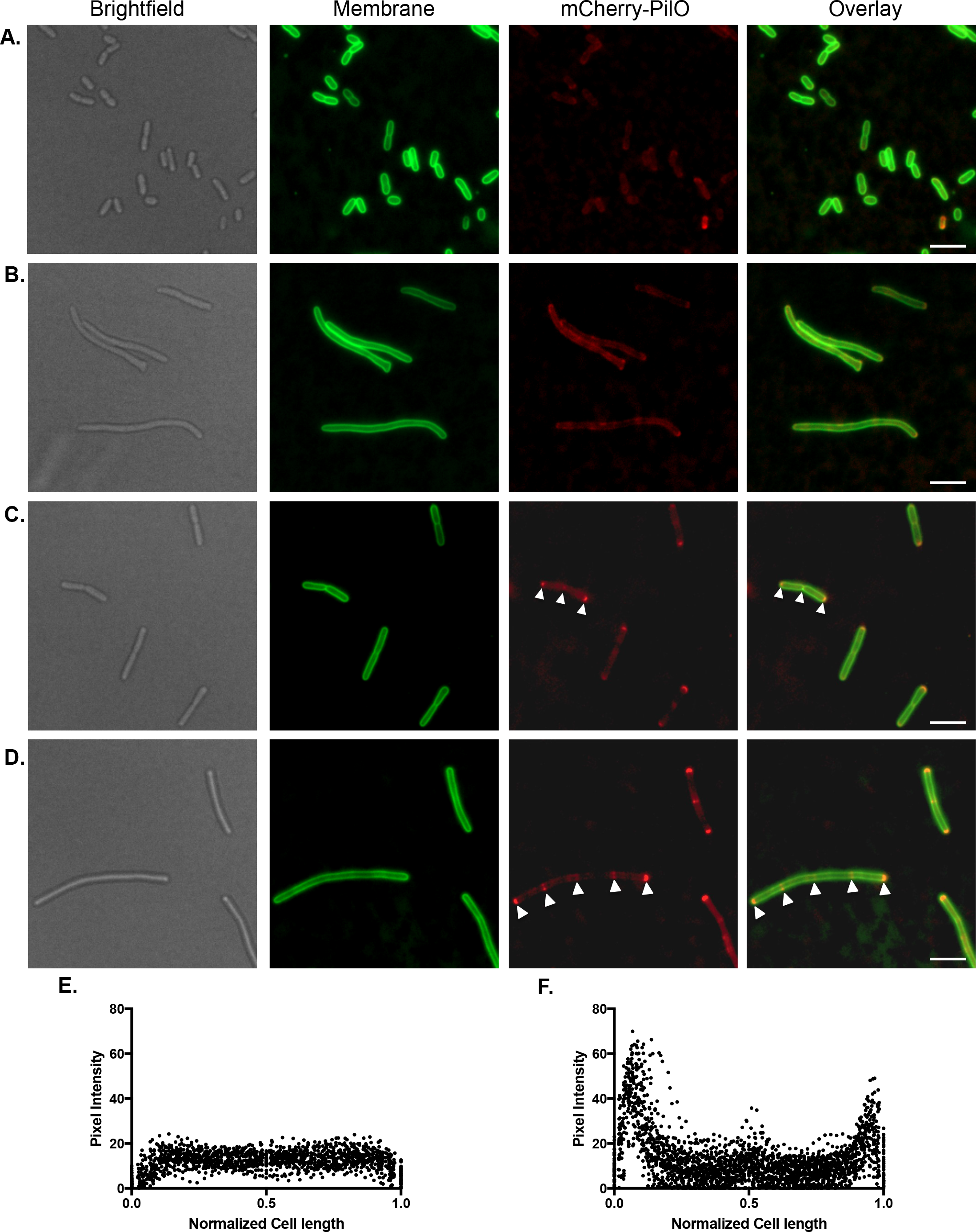
mCherry-PilO is delocalized in a *pocA* mutant. In the absence of the polar organelle coordinating protein, PocA, mCherry-PilO was delocalized in **A**. untreated cells and **B**. cells treated with cefsulodin. When *pocA* was reintroduced *in trans*, mCherry-PilO localization to the poles and septum was recovered (white arrowheads) in **C**. untreated cells, as well as to **D**. the poles and regularly spaced foci in antibiotic-treated cells. The last two panels show quantification of mCherry pixel intensity **E**. in the absence of PocA and **F**. in the pocA-complemented mutant, in 50 untreated cells after normalization of length to 1 as described in the Methods. Scale bar = 3 μm.

**Figure 8.**
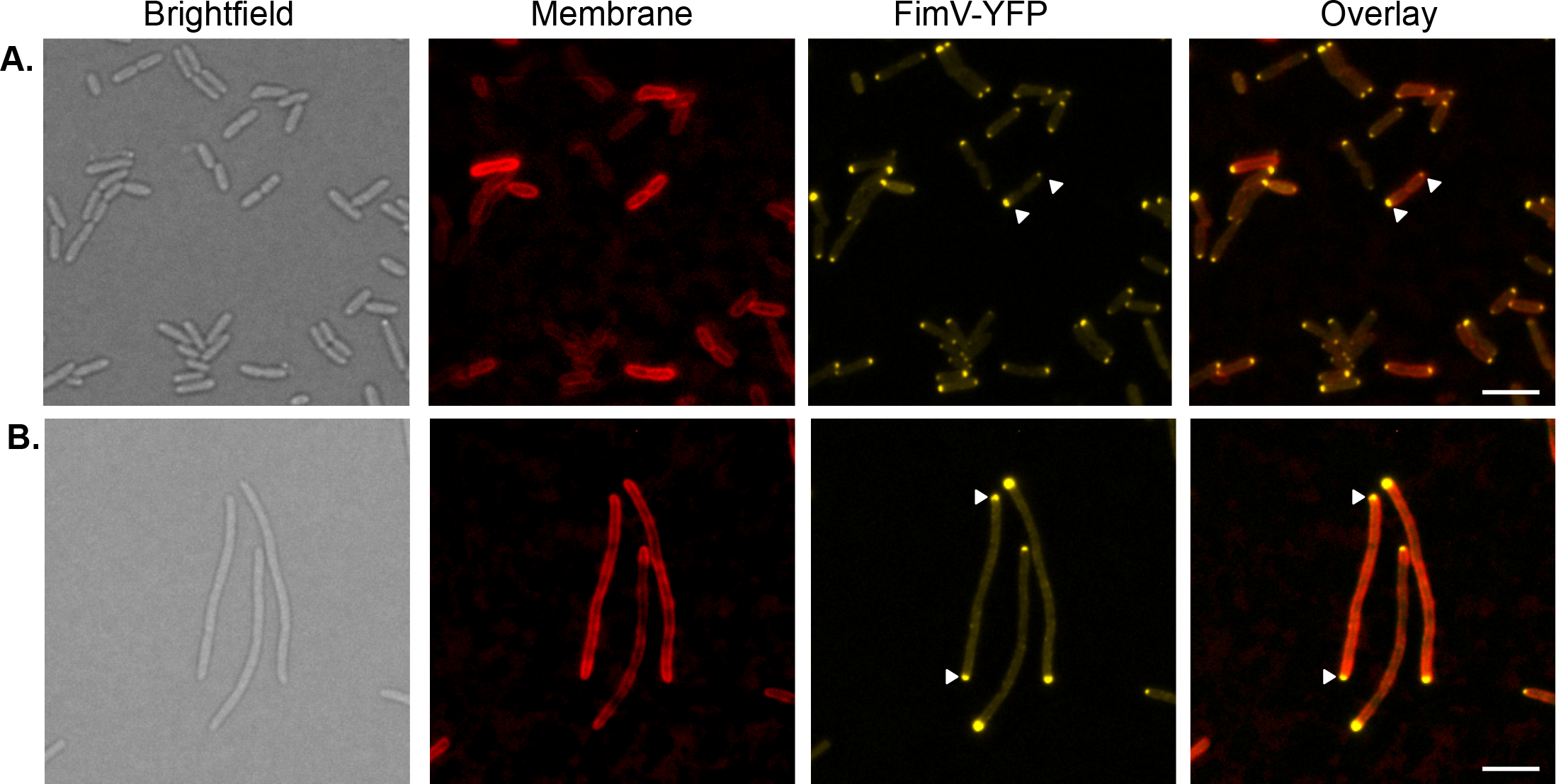
FimV remains polarly localized in the absence of PocA. **A**. FimV-YFP remained localized to the cell poles (arrowheads) in a *pocA* mutant. **B**. FimV-YFP localized to the poles in cefsulodin treated cells. Scale bar = 3 μm.

## DISCUSSION

The T4aP machinery spans all layers of the gram-negative cell envelope at the poles of rod-shaped cells, but how this megadalton protein complex is installed in the absence of dedicated PG hydrolyzing enzymes was unclear. Here we showed that degradation of PG is unnecessary, because components of the *P. aeruginosa* T4P alignment subcomplex and the secretin are recruited to sites of cell division where they can be pre-installed, rather than retrofitted, into nascent poles. These data are consistent with our observation that mutants lacking multiple housekeeping PG hydrolases – which could potentially provide PG lytic activities for retrofitting in the absence of system-specific enzymes – have correctly inserted, functional T4aP machines, as demonstrated by their ability to twitch (**Supplementary Fig. S3**). Instead, at least 3 different mechanisms participate in the process of T4aP polar localization in *P. aeruginosa*, two of which (PilQ and FimV) depend on recognition of PG, as deletion of PilQ’s AMIN domains or FimV’s LysM motif results in delocalization.

Cell poles are characterized by PG that has fewer stem peptides compared to lateral wall PG, due to the activity of N-acetylmuramyl-L-alanine amidases that cleave the amide bond between N-acetylmuramic acid and the stem peptides during daughter cell separation (31–33). This denuded form of PG is the binding target for multiple protein motifs, among them AMIN, SPOR, and LysM (28, 34–36). *P. aeruginosa* and *M. xanthus* PilQ monomers have N-terminal AMIN domains (2 per monomer in *P. aeruginosa* and 3 in *M. xanthus*), which in addition to targeting (37), may anchor secretin complexes to the PG layer to counter the substantial forces generated during twitching motility. *Neisseria gonorrhoeae* and *N. meningitidis* PilQ monomers also contain 2 AMIN domains at their amino termini (11), yet they express peritrichous T4aP. *Neisseria* spp. lack the cytoskeletal element MreB which patterns elongation of rod-shaped bacteria (38), thus have a coccoid morphology. It’s possible that the mechanism of T4aP assembly system recruitment to sites of cell division is similar in *Neisseria* spp., but due to their shape – the equivalent of two closely apposed poles – they appear to have peritrichous T4aP distribution.

In the *P. aeruginosa* T4aP system, at least two different PG-binding proteins were needed for correct positioning. We showed previously (28) that FimV binds PG via its LysM motif, important for optimal secretin assembly. A number of recent studies showed that FimV and related HubP proteins – which contain multiple protein-protein interaction domains in addition to conserved LysM motifs – act as landmarks, responsible for polar localization of other proteins, including those involved in regulation of T4aP function, flagellar placement, and chromosome segregation (39–43). For example, *P. aeruginosa* FimL – a regulatory protein important for T4aP assembly via its regulation of intracellular cAMP levels – interacts with the C-terminal TPR motif of FimV (39, 44). Loss of FimV also results in mislocalization of the core T4aP machinery (**Fig. 6**); thus FimV appears to have both physical and regulatory roles in T4aP assembly and function.

Cowles *et al.* (29) reported that both the flagellum and T4aP were mislocalized in *P. aeruginosa* mutants lacking the polar organelle coordinating (Poc) proteins, but the mechanism remains enigmatic. The T4aP alignment subcomplex was delocalized in a *pocA* mutant (**Fig. 7**) while FimV remained bipolar (**Fig. 8**), implying that they act independently of one another – with the caveat that the FimV fusion used here was non-functional in twitching motility. PocAB-TonB3 are homologous to the ExbBD-TonB complex in *E. coli* (29) which energizes siderophore uptake across the OM via direct interactions with the ‘Ton box’ on one beta strand of TonB-dependent receptors. The N0 domains of secretin monomers are structurally similar (2.3 Å root mean square deviation over 65 residues) to the signalling domain of FpvA, a TonB-dependent siderophore receptor in *P. aeruginosa* (45). This similarity hints at potential interactions between the TonB3 component of the Poc complex and the N0 domain of PilQ monomers. We noted that although deletion of *pocA* had no significant effect on cell morphology, complementation with a plasmid-borne copy of the gene increased average cell length by ∼50% (**Supplementary Fig. S4**), suggesting that the Poc system might influence the timing of cell division or affect PG remodelling. Subtle structural differences in PG architecture in the absence of PocA could impact binding of proteins with PG targeting motifs.

Mid-cell recruitment of mCherry-PilO was dependent on the secretin monomer PilQ (**Fig. 3CD**) but deletion of the pilotin PilF had no effect (**Fig. 5**), showing that recruitment and confinement of PilMNOPQ to cell division sites occur prior to secretin oligomerization. Prior studies of *M. xanthus* T4aP assembly system formation suggested that PilQ multimerization was essential for the recruitment of additional T4aP assembly components, because loss of Tgl – the *M. xanthus* PilF homolog – abrogated polar localization of PilQ and other T4aP proteins. These data supported an ‘outside in’ model for *M. xanthus* T4aP system formation, dependent on PilQ oligomerization (26). Interestingly, *M. xanthus* PilQ was suggested in that study to localize to the septum either “late during cell division, or immediately after”. These data are consistent with the pattern of PilQ localization in *P. aeruginosa:* however, the pilotin proteins in *M. xanthus* and *P. aeruginosa* appear to play somewhat different roles.

Together, the data led us to propose a pathway for T4aP assembly machinery integration into the *P. aeruginosa* cell envelope in the absence of dedicated PG hydrolases. We suggest that PilMNOPQ are co-translated, forming an IM-bound subcomplex that diffuses laterally until the AMIN domains of PilQ monomers bind to septal PG. FimV, which has the capacity to interact with septal PG via its Lys motif, may also interact with components of the subcomplex to retain them at cell division sites. The next steps are still unclear, as the oligomerization pathway for PilQ in the OM remains unknown. We previously proposed (9) that co-translocation of the lipoprotein PilF and PilQ monomers by the Lol system might promote PilQ insertion into the OM, where it oligomerizes. In this new model, we suggest that PilF is translocated independently by the Lol system to the OM, where it then scans for, binds to, and promotes translocation of PilQ monomers from the inner to the OM during invagination in the final stages of cell division.

Two lines of evidence support this new hypothesis. First, in *M. xanthus*, coincubation of *pilQ* and *tgl (pilF*) mutants, neither of which are able to twitch, allowed for <u>inter</u>cellular transfer of Tgl from the *pilQ* mutant to the *tgl* mutant and oligomerization of the latter’s PilQ monomers, restoring piliation and motility (46). Thus, Tgl already present in the OM of the *pilQ* mutant could promote insertion and oligomerization of PilQ monomers in the *tgl* mutant. Second, the ability of OM lipoproteins to interact with inner membrane proteins across the periplasm is now well established, as exemplified by the *E. coli* OM lipoproteins, LpoA and LpoB (47, 48). These proteins are essential for stimulating the activities of the inner membrane PG synthases PBP1a and PBP1b, respectively, through direct interactions with regulatory domains of those enzymes. Once PilQ monomers reach the OM with the assistance of PilF, they may first form conformational intermediates prior to maturation into stable, SDS-resistant oligomers. In *N. gonorrhoeae*, certain point mutations in the secretin domain were associated with formation of immature, leaky oligomers, rendering the cells more susceptible to beta lactam antibiotics (49).

The above scenario for PilQ translocation suggests that the ∼ 2 nm porosity of PG (50) would not pose a physical barrier to secretin assembly, as individual monomers (∼77 kDa) are small enough to pass through its natural gaps. Retention of PilQ AMIN interactions with septal PG during translocation and insertion of the C-terminal secretin domain in the OM would ultimately place the AMIN domains ‘under’ the PG layer, where they would be ideally positioned to counteract retraction forces imposed during twitching motility. Because polar PG is less likely to be remodelled due to its limited side chain content, the secretin would remain stably fixed to the cell wall. Studies of the half-life of a single bacterial cell pole show that it can be hundreds of generations old (51); thus, it is perhaps not surprising that secretin complexes are highly resistant to denaturation, allowing them to last as long as the poles in which they are embedded.

Among the questions remaining is whether the interaction interface between PilP and PilQ is altered during translocation of PilQ monomers to the OM. We established that mid-cell localization of mCherry-PilO depends on PilP (**Fig. 3EF**), which interacts with the N0 domain of PilQ monomers via its C-terminal HR domain (11, 20). PilP also interacts with PilNO heterodimers via its long unstructured N-terminus (17). This flexible tether might allow the C-terminal domain of PilP to remain connected to the N0 domain of PilQ as secretin monomers transit from the inner to OM. Alternatively, PilP may be repositioned relative to PilQ when the latter translocates to the OM. In the evolutionarily related T2SS, multiple interaction interfaces between the HR domain of GspC (PilP) and N0 of GspD (PilQ) have been identified through a combination of structural, biochemical, and functional studies (45, 52, 53). These various interfaces – some of which are mutually exclusive – were attributed to differences in experimental setup, but could also reflect capture of different interaction states pre- and post-secretin oligomerization. Studies to address these possibilities are currently underway.

## MATERIALS & METHODS

### Strains, media and growth conditions

Bacterial strains, plasmids, and primers used in this study are listed in Tables 1 and 2. *Escherichia coli* and *P. aeruginosa* were grown at 37°C in LB Lennox broth or on LB 1.5% agar plates supplemented with antibiotics. The antibiotics and their respective concentrations were: ampicillin (Ap), 100 μg/mL; carbenicillin (Cb), 200 μg/mL; gentamicin (Gm), 15 μg/mL for *E. coli* strain and 30 μg/mL for *P. aeruginosa* strains, unless otherwise stated. Plasmids were transformed by heat shock into chemically competent *E. coli* cells or by electroporation into *P. aeruginosa* suspended in sterile water. All constructs were verified by DNA sequencing (MOBIX Lab, McMaster University).

**Table 1.**
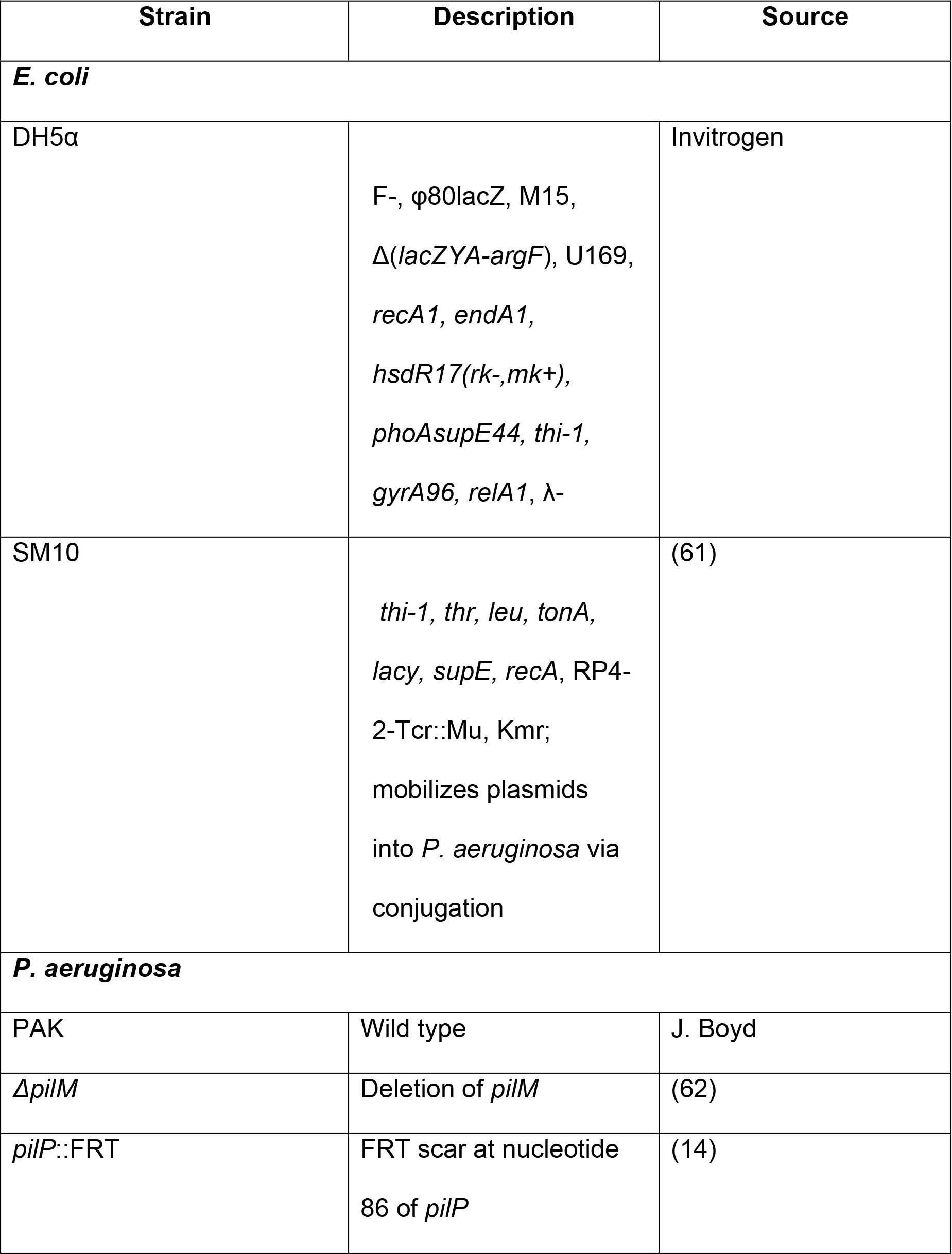
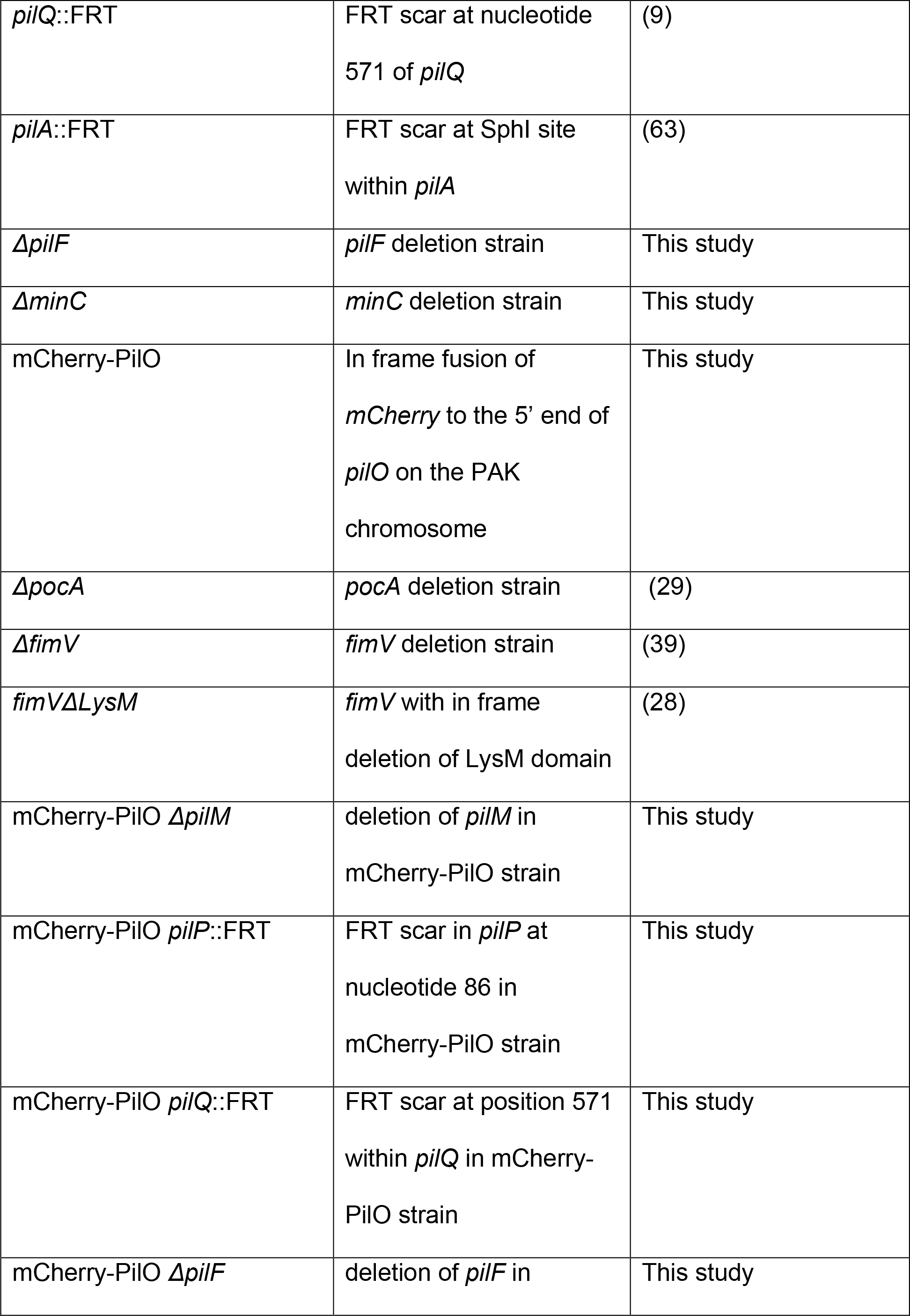
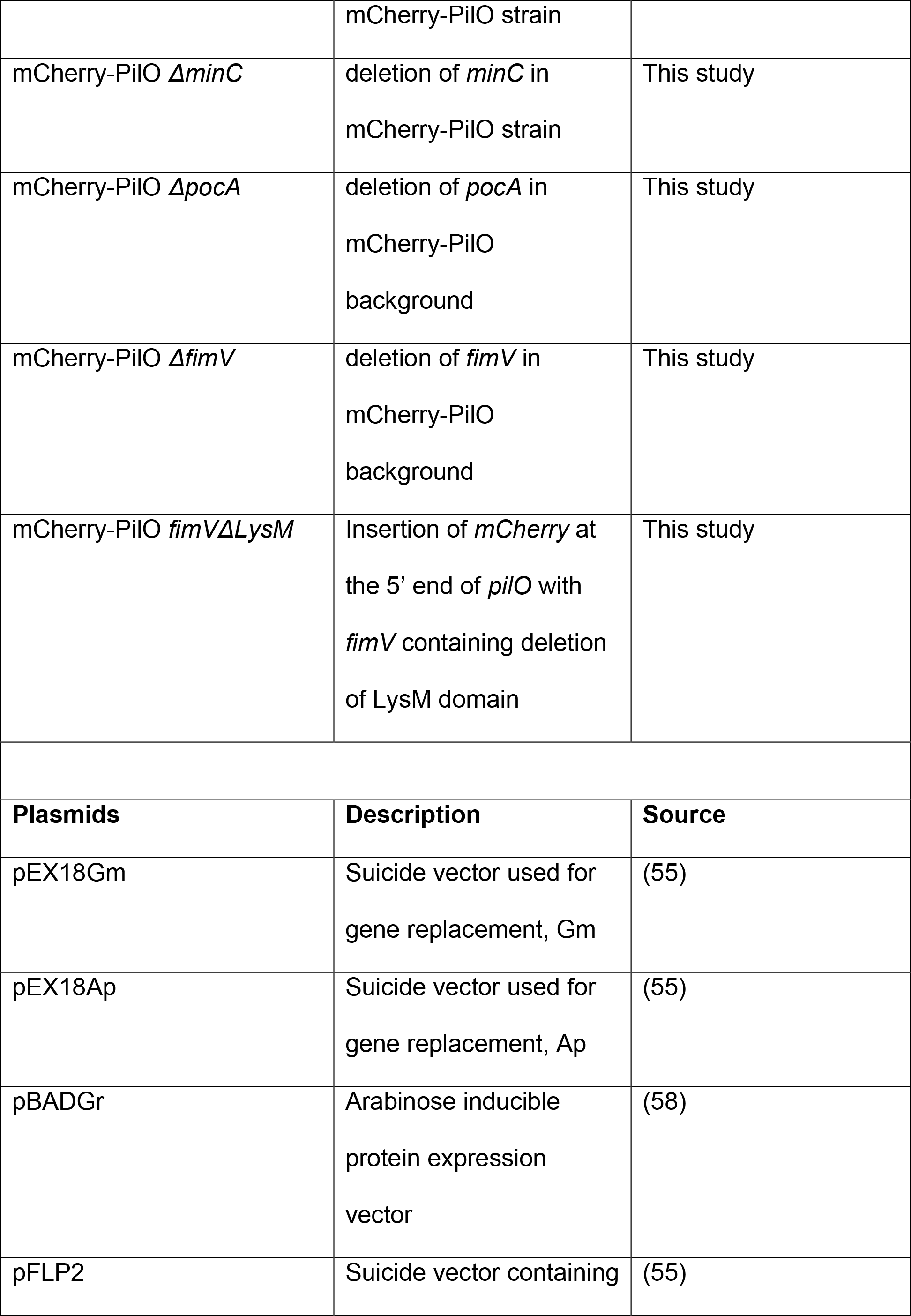
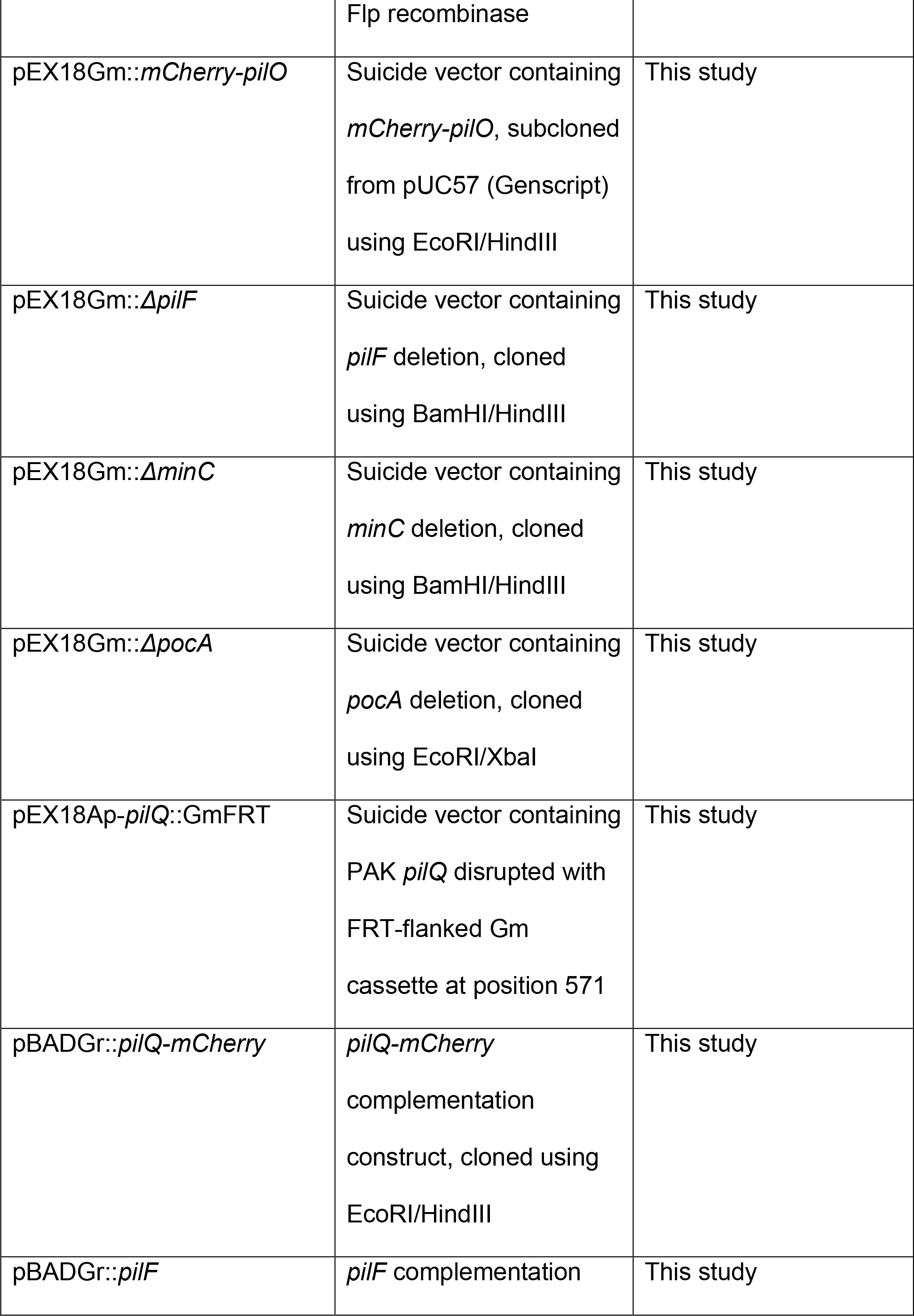
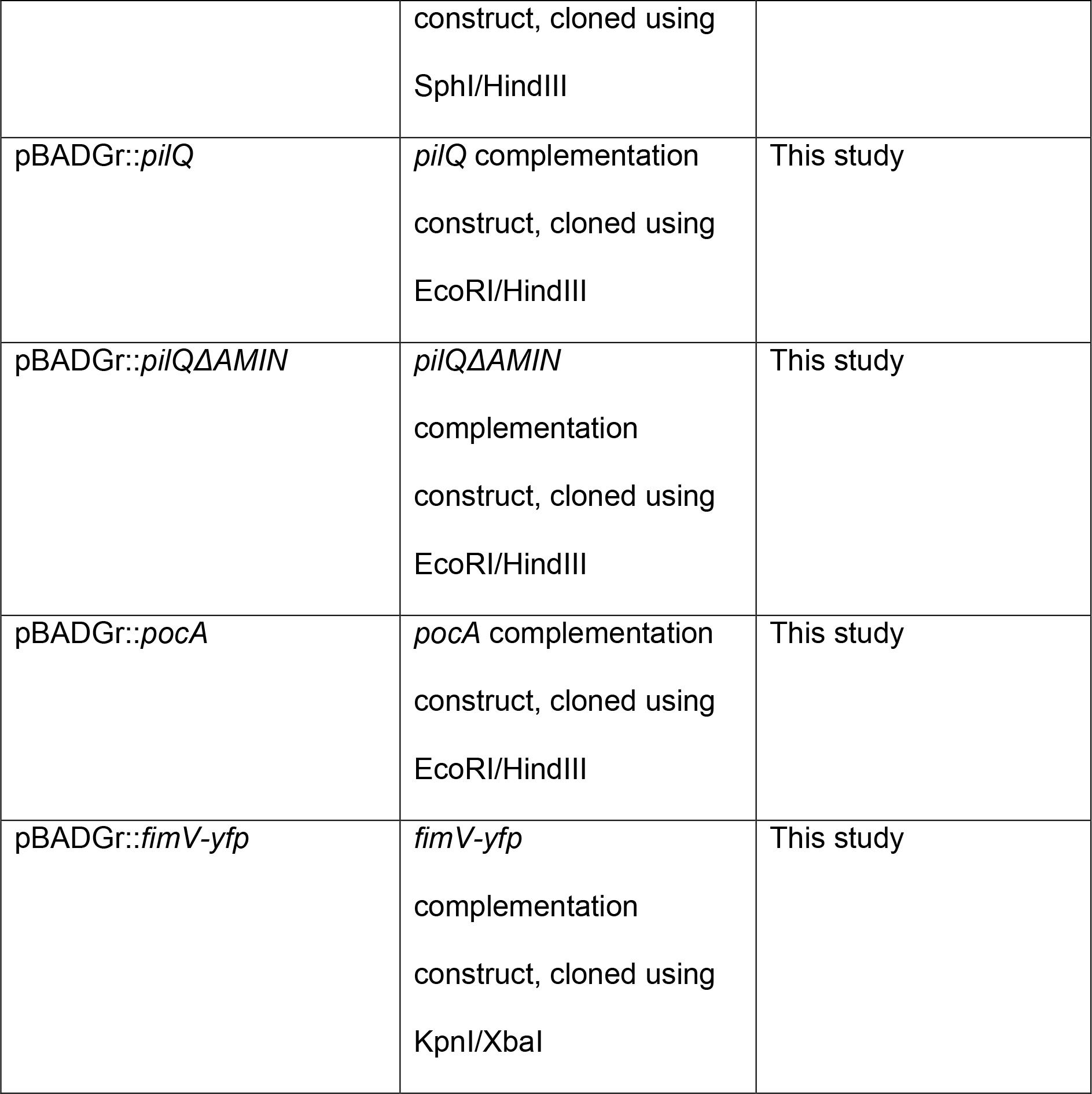
Bacterial strains and plasmids

| Strain | Description | Source |
| --- | --- | --- |
| ***E. coli*** |  |  |
| DH5α | F<sup>-</sup>, ϕ80lacZ, M15,<br>Δ(*lacZYA-argF*), U169,<br>*recA1*, *endA1*,<br>*hsdR17(rk<sup>-</sup>,mk<sup>+</sup>)*,<br>*phoA*supE44, *thi-1*,<br>*gyrA96*, *relA1*, λ<sup>-</sup> | Invitrogen |
| SM10 | *thi-1*, *thr*, *leu*, *tonA*,<br>*lacy*, *supE*, *recA*, RP4-2-Tcr::Mu, Kmr;<br>mobilizes plasmids<br>into *P. aeruginosa* via<br>conjugation | (61) |
| ***P. aeruginosa*** |  |  |
| PAK | Wild type | J. Boyd |
| Δ*pilM* | Deletion of *pilM* | (62) |
| *pilP*::FRT | FRT scar at nucleotide<br>86 of *pilP* | (14) |
| *pilQ*::FRT | FRT scar at nucleotide 571 of *pilQ* | (9) |
| *pilA*::FRT | FRT scar at SphI site within *pilA* | (63) |
| $\Delta pilF$ | *pilF* deletion strain | This study |
| $\Delta minC$ | *minC* deletion strain | This study |
| mCherry-PilO | In frame fusion of *mCherry* to the 5' end of *pilO* on the PAK chromosome | This study |
| $\Delta pocA$ | *pocA* deletion strain | (29) |
| $\Delta fimV$ | *fimV* deletion strain | (39) |
| *fimV*ΔLysM | *fimV* with in frame deletion of LysM domain | (28) |
| mCherry-PilO $\Delta pilM$ | deletion of *pilM* in mCherry-PilO strain | This study |
| mCherry-PilO *pilP*::FRT | FRT scar in *pilP* at nucleotide 86 in mCherry-PilO strain | This study |
| mCherry-PilO *pilQ*::FRT | FRT scar at position 571 within *pilQ* in mCherry-PilO strain | This study |
| mCherry-PilO $\Delta pilF$ | deletion of *pilF* in | This study |
|  | mCherry-PilO strain |  |
| mCherry-PilO $\Delta minC$ | deletion of *minC* in mCherry-PilO strain | This study |
| mCherry-PilO $\Delta pocA$ | deletion of *pocA* in mCherry-PilO background | This study |
| mCherry-PilO $\Delta fimV$ | deletion of *fimV* in mCherry-PilO background | This study |
| mCherry-PilO *fimV*ΔLysM | Insertion of *mCherry* at the 5' end of *pilO* with *fimV* containing deletion of LysM domain | This study |
| **Plasmids** | **Description** | **Source** |
| pEX18Gm | Suicide vector used for gene replacement, Gm | (55) |
| pEX18Ap | Suicide vector used for gene replacement, Ap | (55) |
| pBADGr | Arabinose inducible protein expression vector | (58) |
| pFLP2 | Suicide vector containing | (55) |
|  | Flp recombinase |  |
| pEX18Gm::*mCherry-pilO* | Suicide vector containing *mCherry-pilO*, subcloned from pUC57 (Genscript) using EcoRI/HindIII | This study |
| pEX18Gm::*ΔpilF* | Suicide vector containing *pilF* deletion, cloned using BamHI/HindIII | This study |
| pEX18Gm::*ΔminC* | Suicide vector containing *minC* deletion, cloned using BamHI/HindIII | This study |
| pEX18Gm::*ΔpocA* | Suicide vector containing *pocA* deletion, cloned using EcoRI/XbaI | This study |
| pEX18Ap-*pilQ*::GmFRT | Suicide vector containing PAK *pilQ* disrupted with FRT-flanked Gm cassette at position 571 | This study |
| pBADGr::*pilQ-mCherry* | *pilQ-mCherry* complementation construct, cloned using EcoRI/HindIII | This study |
| pBADGr::*pilF* | *pilF* complementation | This study |
|  | construct, cloned using<br>SphI/HindIII |  |
| pBADGr::*pilQ* | *pilQ* complementation<br>construct, cloned using<br>EcoRI/HindIII | This study |
| pBADGr::*pilQ*Δ*AMIN* | *pilQ*Δ*AMIN*<br>complementation<br>construct, cloned using<br>EcoRI/HindIII | This study |
| pBADGr::*pocA* | *pocA* complementation<br>construct, cloned using<br>EcoRI/HindIII | This study |
| pBADGr::*fimV-yfp* | *fimV-yfp*<br>complementation<br>construct, cloned using<br>KpnI/XbaI | This study |

**Table 2.**
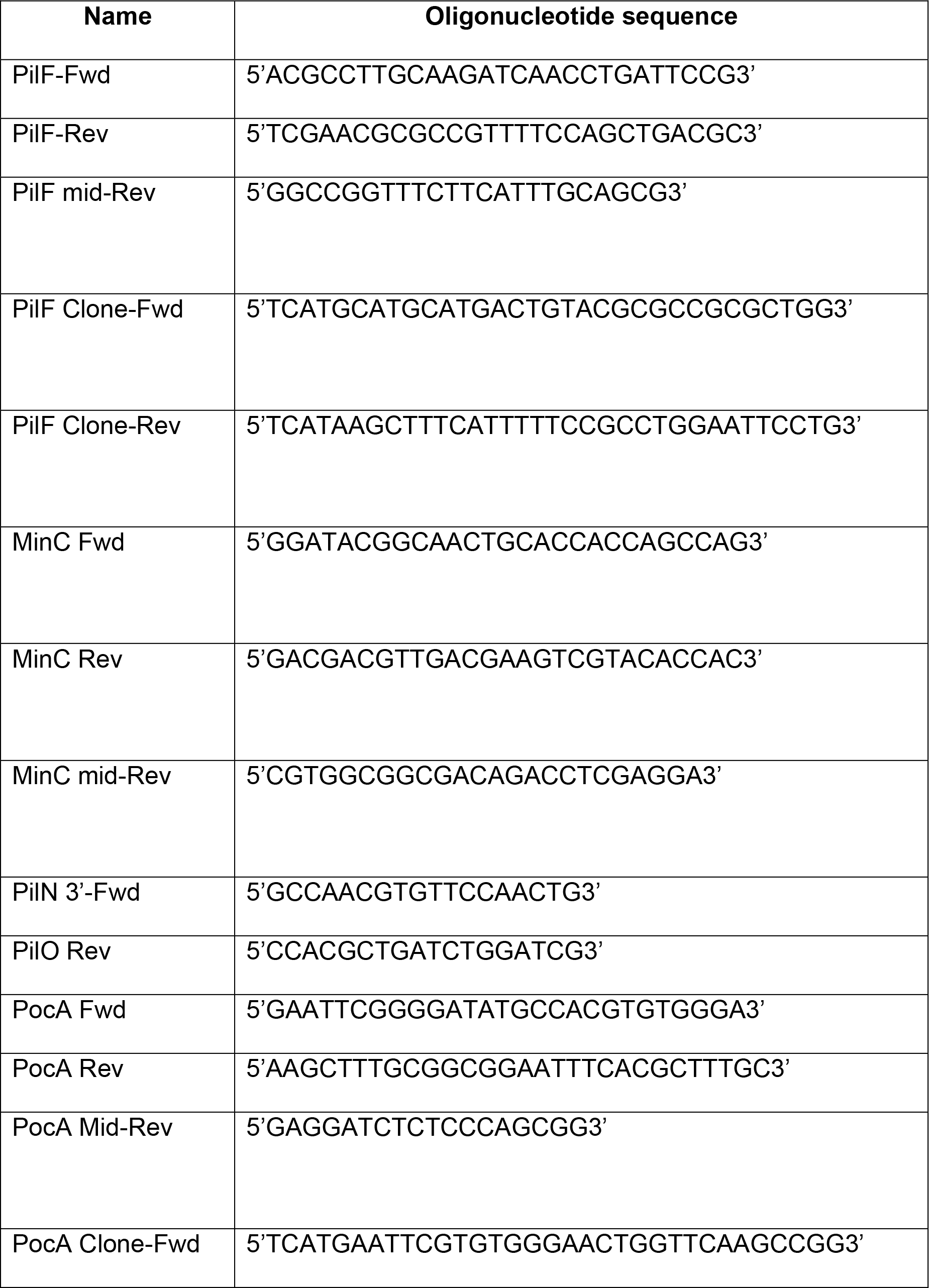
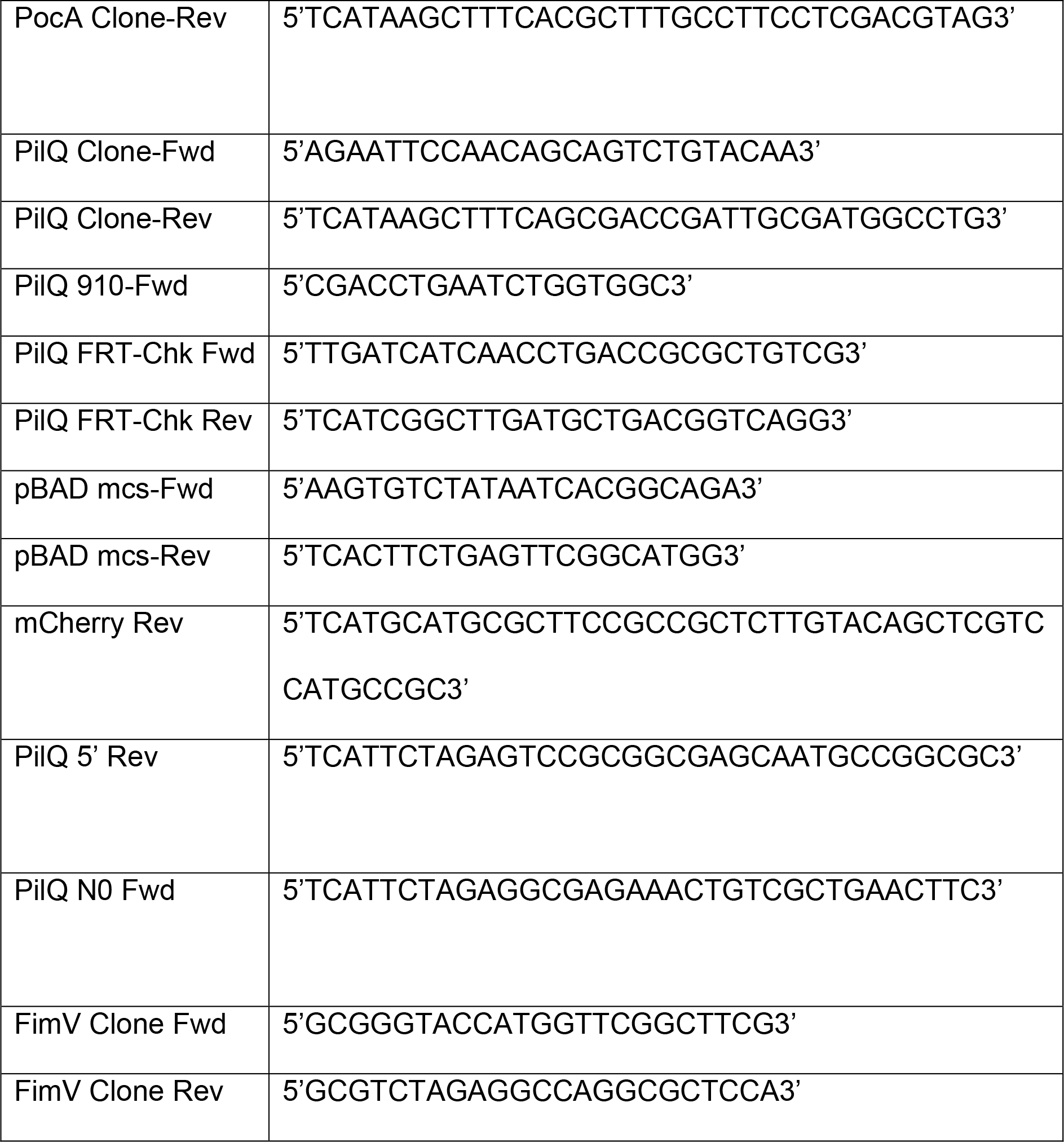
Primer sequences

### Generation of *P. aeruginosa* mutants

Mutants were made using previously described methods (54). Deletion constructs for generation of *minC* and *pilF* mutants were designed to include 500 nucleotides up and downstream of the gene to be deleted, as well as the first and last 50 nucleotides of the gene. Constructs were synthesized and cloned into pUC57 (Genscript, Piscataway, NJ). Deletion inserts in pUC57 were subcloned into the pEX18Gm suicide vector (55) using the 5’ BamHI and the 3’ HindIII restriction sites, and verified by DNA sequencing. After verification, suicide vectors containing the mutant genes (*pilF* or *minC*) were transformed into *E. coli* SM10 cells. Plasmids were then transferred by conjugation in a 1:9 ratio of *P. aeruginosa* to *E. coli.* The mixed culture was pelleted for 1 min at 2292 × *g*, and the pellet was resuspended in a 100 μL aliquot of LB, spot-plated on LB agar, and incubated overnight at 37 °C. The mating mixture was removed from the LB agar using a sterile toothpick, resuspended in 1 mL of LB and the *E. coli* SM10 donor was counterselected by plating the mixture on *Pseudomonas* isolation agar (PIA; Difco) containing Gm (100 μg/ml). Gm-resistant *P. aeruginosa* colonies were streaked on LB no salt plates containing sucrose (1% (w/v) bacto-tryptone, 5% (w/v) sucrose, 1.5% agar, 0.5% (w/v) bacto-yeast extract) and incubated for 16 h at 30 °C. Select colonies were then subcultured on LB agar and LB agar plus Gm, and Gm-sensitive colonies were screened by PCR using appropriate primers (Table 2) to confirm integration. Amplicons from colonies with the desired PCR profile were sequenced to confirm incorporation of the mutation, and the *pilF* mutant was analyzed by western blot analysis using an α-PilF antibody to verify loss of the gene product.

A *pilQ* KO construct containing an FRT-flanked Gm cassette was designed previously (9). This construct was integrated into the PAK chromosome using a Flp-FRT recombination system (55) and selected for using the methods described above with the following modifications: following growth on sucrose, colonies were cultured on LB agar, LB agar with Gm and LB agar plus Cb. Colonies that had undergone double recombination, integrating pilQ::GmFRT into the chromosome without pEX18Ap (Cb sensitive colonies), were selected. The Gm cassette was removed by Flp recombinase-catalyzed excision, by conjugally transferring the Flp-expressing pFLP2 from SM10 cells into PAK cells carrying the GmFRT-disrupted *pilQ* gene. SM10 cells were counterselected on PIA containing Cb (200 μg/ml). pFLP2 was removed from *P. aeruginosa* by streaking Cb-resistant colonies on LB agar, no salt, containing 5% (w/v) sucrose for 16 h at 30 °C. Colonies were then cultured on LB agar and LB-agar containing Gm or Cb. Cb and Gm sensitive colonies were screened by PCR using the PilQ FRT-Chk primers listed in Table 2 to confirm the retention of the FRT scar. The *pilQ* gene of mutant strains were sequenced to confirm correct incorporation of the mutation, and analyzed by western blot with α-PilQ sera to confirm loss of the gene product.

The *pocA* deletion construct was created previously in the pEX18Tc suicide vector (29). The *pocA* deletion insert was subcloned into pEX18Gm using EcoRI and Xbal, and constructs verified by DNA sequencing. After verification, the suicide vector containing the *pocA* deletion construct transformed into *E. coli* SM10 cells and the mutation introduced into *P. aeruginosa* as described above.

### Generation of a chromosomally encoded mCherry-PilO fusion

A construct encoding an fusion of mCherry (56) to the N-terminus of PilO, plus 700 nucleotides upstream of *pilO* and 700 nucleotides downstream of *pilO*, was synthesized and cloned into pUC57 (Genscript, Piscataway, NJ). The mCherry-PilO-encoding insert was subcloned into the suicide vector pEX18Gm using the 5’ EcoRI and the 3’ Hindlll restriction sites, constructs verified by DNA sequencing, and introduced into chemically competent *E. coli* SM10 by heat shock. The plasmid was then transferred by conjugation in a 1:9 ratio of *P. aeruginosa* (wild type or various T4aP mutants) to *E. coli* as described above. Following counter selection and curing of merodiploids by sucrose selection, integration of the mCherry-PilO fusion in Gm-sensitive colonies was confirmed by PCR and DNA sequencing (Table 2). mCherry-PilO expressing strains were analyzed using western blot analysis and twitching assays to verify expression of an intact fluorescent fusion protein and – in the wild type – function of the T4aP machinery, respectively.

### Generation of complementation constructs

The *pilQ, pocA* and *pilF* genes were amplified by PCR using PAK chromosomal DNA as the template (Table 2) and cloned into pBADGr (Table 1). The digested DNA was purified and ligated into pBADGr with T4 DNA ligase, according to the manufacturer’s instructions. Ligation mixtures were then transformed into *E. coli* DH5a cells and grown overnight at 37 °C on LB agar supplemented with the appropriate antibiotics.

A construct encoding a truncated version of PilQ lacking its AMIN domains was previously generated in the PAO1 strain (9). The same boundaries were used here to generate a truncated version of PAK PilQ. The PilQ Clone Fwd and PilQ 5’ Rev primers were used to amplify the 5’ end of the gene, excluding the first AMIN domain, and the PilQ N0 Fwd and PilQ Clone Rev primers (Table 2) were used to amplify the remainder of the gene beginning at the N0 domain and excluding the second AMIN domain. Purified amplicons were digested with Xbal and ligated into pBADGr with T4 DNA Ligase. The ligation product and pBADGr were then digested with EcoRI and Hindlll and ligated following the same protocol. The fidelity of the pBADGr-*pil*Q construct was confirmed by sequencing using the pBADGr mcs Fwd and Rev primers (Table 2).

Since PilQ has a cleavable N-terminal signal sequence, and a C-terminus that is exposed to the extracellular environment (9, 57), we designed a construct encoding an internal fusion of mCherry to PilQ (between the first and second AMIN domains, residues 132 and 133), which was synthesized and subcloned into pUC57 (Genscript). The PilQ-mCherry-encoding insert was subcloned into the complementation vector pBADGr (58) using the 5’ EcoRI and the 3’ HindIII restriction sites and verified by DNA sequencing. The construct was then introduced into PAK *pil*Q::FRT cells by electroporation. Expression of the PilQ-mCherry fusion was verified using western blot analysis with α-PilQ and α-mCherry antibodies and function was verified using twitching assays (59).

The *yfp* gene encoding Yellow Fluorescent Protein (YFP) was cloned into the HindIII restriction site of pBADGr. Correct insertion of the pBADGr-yfp construct was confirmed by DNA sequencing using mcs Fwd/Rev primers (Table 2). A version of *fimV* lacking its stop codon was amplified using the FimV Clone Fwd/Rev primers (Table 2) and cloned into pBADGr in frame with the *yfp* gene by digesting both the vector and insert with KpnI and XbaI restriction enzymes and ligating with T4 DNA ligase. Successful ligation of the insert was confirmed using the pBADGr multiple cloning site flanking primers (Table 2) and DNA sequencing. The construct was then electroporated into *ΔfimV* and *ΔpocA* cells. A stable fusion of YFP to the C-terminus of FimV was confirmed by western blotting with α-AFP and α-FimV antibodies. This fusion is considered non-functional as it failed to restore twitching in the *fimV* background.

### Twitching motility assays

Twitching motility assays were performed as previously described (59). Single colonies were stab inoculated to the bottom of a 1 *%* LB agar plate. Plates were incubated for 24 h at 30 °C. Following incubation, the agar was removed and the adherent bacteria stained with 1 % (w/v) crystal violet for 30 min, followed by washing with water to remove unbound dye. Twitching zones were photographed and their areas measured using Fiji (60). All experiments were performed in triplicate with at least three independent replicates.

### Analysis of whole cell lysates by western blot

Cultures were grown overnight at 37 °C in LB supplemented with appropriate antibiotics and diluted to an OD_600_ of 0.6. A 1 mL aliquot of cells was collected by centrifugation at 2292 × *g* for 1 min. and the cell pellet was resuspended in 100 μl of SDS sample buffer (80 mM Tris (pH 6.8), 5.3% (v/v) 2-mercaptoethanol, 10% (v/v) glycerol, and 0.02% (w/v) bromophenol blue, and 2% (w/v) SDS) and boiled for 10 min.

Equal volumes of whole cell lysates were separated by 12.5% SDS-PAGE at 80-150 V and transferred to nitrocellulose membranes for 1 h at 225 mA. Membranes were blocked using 5% (w/v) low fat skim milk powder in phosphate buffered saline solution (PBS) at pH 7.4 for 1 h at room temperature, followed by incubation with the appropriate polyclonal (for PilF, PilM, PilO, PilP, PilQ, FimV) or monoclonal (for mCherry) antisera for 1 h at room temperature. Membranes were washed twice in 10 mL of PBS for 5 min then incubated in either goat-anti-rabbit or goat-anti-mouse IgG-alkaline phosphatase conjugated secondary antibody (Bio-Rad) at a dilution of 1:3000 for 1 h at room temperature. Membranes were washed twice in PBS for 5 min, and developed in a solution containing 100 μL nitro-blue tetrazolium (NBT) and 100 μL 5-bromo-4-chloro-3-indolyl phosphate (BCIP) in 10 mL of alkaline phosphatase buffer (100 mM NaCl, 5 mM MgCl_2_, 100 mM Tris, pH 9.5).

### Cell preparation for transmission electron microscopy

Cells from a 1 mL overnight culture in LB were pelleted at 2292 × *g* and resuspended in a 1 mL solution of 2% (w/v) gluteraldehyde in 0.1M sodium cacodylate buffer at a pH of 7.4. Cells were fixed in solution for 2 h at 4 °C, and washed twice with cold sodium cacodylate buffer (pH 7.4). Samples were then treated with 1% osmium tetroxide in 0.1M sodium cacodylate for 1 h and stained with 2% uranyl acetate overnight. After staining, cells were dehydrated gradually with ethanol, treated with propylene oxide, embedded in Spurr’s resin, and polymerized at 60 °C overnight. The polymerized samples were sectioned using a Leica UCT ultramicrotome and poststained with 2% uranyl acetate and lead citrate. The prepared specimens were examined in McMaster University’s Electron Microscopy Facility using the JEOL JEM 1200 EX TEMSCAN microscope (JEOL, Peabody, MA, USA) operating at an accelerating voltage of 80kV. Images were acquired with an AMT 4-megapixel digital camera (Advanced Microscopy Techniques, Woburn, MA).

### Fluorescence Microscopy

Strains were incubated overnight at 37 °C in 5 mL of LB supplemented with the appropriate antibiotics. One mL of cells were pelleted at 2292 × g, and resuspended in 1 mL of sterile water containing 10 μg/ml of either Fm1-43 Fx (for cells expressing mCherry) or Fm1-64 Fx (for cells expressing YFP) membrane stain (Invitrogen). Cells were mixed with dye by gentle pipetting, and pelleted at 2292 × g. Pelleted cells were then stab-inoculated using a pipette tip to the coverslip-media interface at the bottom of an 8-well microscope glass-coverslip slide containing 200 μl 1% LB agarose per well (Lab-Tek). Cells were then incubated for 75 min at 37 °C prior to imaging. In filamentation experiments, 1 mL of cells grown overnight were sub-cultured into 4 mL of LB containing 40 μg/ml of cefsulodin and incubated for 3 h at 37 °C. Cells were then stained and mounted for microscopy using the above protocol. Cells were imaged using an EVOS Fl-auto microscope in the McMaster Biophotonics Facility. Images were acquired using a 60x oil immersion objective using a Texas Red filter, YFP filter, or transmitted white light (no filter). Fiji (60) was used to adjust image brightness and contrast, and to overlay brightfield and fluorescent images.

### Quantification of mCherry-PilO fluorescence

Fluorescence was quantified using Fiji software (60). Each field in the Texas Red (for mCherry) and YFP (for Fm1-43 Fx) filter sets were overlaid, and used with the micrograph from the YFP filter for image analysis. A 250 μm^2^ grid was added to each field for precise documentation of the quantified cells. Moving systematically between squares in the grid, cells that fulfilled the following criteria were selected: 1) not contacting other cells; 2) normal rod morphology; and 3) a stable mCherry-PilO fusion (diffuse cytoplasmic fluorescence implies cleavage of the tag in that cell). Cell lengths and pixel intensities were measured from one cell pole to the other and normalized to 1 (i.e., each data point associated with length divided by the total length of the cell to yield normalized 0 to 1 scale). Pixel intensity measurements in the YFP field (cell outline) were then subtracted from the overlay field, producing mCherry-PilO pixel intensity measurements alone in each cell on a scale from 0 to 1. From the data, an XY scatter plot was generated to show the distribution of mCherry-PilO fluorescence over cell length.

## ACKNOWLEDGEMENTS

We thank Dr. Zemer Gitai for the *pocA* deletion construct, and Dr. Ray Truant and McMaster Biophotonics for access to the EVOS microscope, and the McMaster Electron Microscopy facility for assistance with TEM.

